# Surface electric fields increase human osteoclast resorption through improved wettability on carbonate–incorporated apatite

**DOI:** 10.1101/2021.07.28.454035

**Authors:** Leire Bergara Muguruza, Keijo Mäkelä, Tommi Yrjälä, Jukka Salonen, Kimihiro Yamashita, Miho Nakamura

**Affiliations:** Medicity Research Laboratory, Faculty of Medicine, University of Turku, Tykistökatu 6, 20520, Turku, Finland; Turku University Hospital, University of Turku, Luolavuorentie 2, 20700 Turku, Finland; Department of Anesthesia and Intensive Care, University of Turku, Luolavuorentie 2, 20700 Turku, Finland; Graduate School of Medical and Dental Science, Tokyo Medical and Dental University, 1–5–45, Yushima, Bunkyo–ku, Tokyo, 113–8510, Japan; Institute of Biomaterials and Bioengineering, Tokyo Medical and Dental University, 2–3–10 Kanda–Surugadai, Chiyoda, Tokyo 1010062 Japan

## Abstract

Osteoclast-mediated bioresorption can be of an efficient means of incorporating the dissolution of biomaterials in the bone remodeling process. Because of compositionally and structurally close resemblance of biomaterials with the natural mineral phases of the bone matrix, synthetic carbonate-substituted hydroxyapatite (CA) is considered as an ideal clinical biomaterial. The present study therefore investigated the effects of electrical polarization on the surface characteristics and interactions with human osteoclasts of hydroxyapatite (HA) and CA. Electrical polarization was found to improve the surface wettability of these materials by increasing the surface free energy, and this effect was maintained for one month. Analyses of human osteoclast cultures established that CA subjected to a polarization treatment accelerated osteoclast resorption but did not affect the early differentiation phase or the adherent morphology of the osteoclasts as evaluated by staining. These data suggest that the surface characteristics of the CA promoted osteoclast resorption. The results of this work are expected to contribute to the design of cell-mediated biomaterials that can be resorbed by osteoclasts after fulfilling their primary function as a scaffold for bone regeneration.

## Introduction

Cell-mediated bioresorption is a biological process in which biomaterials are resorbed by cells and thereby either partially or completely disappear from implantation sites over a period of time ^1^. In the case of an ideal bioresorbable biomaterial intended for bone regeneration, no foreign material will remain after bone restoration and the load-bearing capacity at the restored site will be similar to that of the natural bone tissue ^1^. Hydroxyapatite (HA) and β-tricalcium phosphate (β-TCP) have excellent biocompatibility and osteoconductivity, both of which are required for orthopaedic biomaterials, and so are frequently used in clinical work. HA is a poorly resorvable biomaterial while β–TCP is soluble *in vivo*. The degradation of β-TCP thus proceeds via solution-mediated chemical dissolution, such that this material will dissolve under physiological conditions. Cell-mediated bioresorption is a new technique that is advantageous because it allows for the dissolution of biomaterials after the bone remodeling process in conjunction with bone resorption and formation. Because osteoclasts are responsible for bone resorption, the development of osteoclast-mediated bioresorbable biomaterials is imperative for bone regeneration in vivo. Interestingly, the incorporation of carbonate ions within the HA crystal structure has been experimentally validated to increase osteoclast differentiation and resorption ^2, 3^, even though stoichiometric HA cannot be resorbed by osteoclasts. Carbonate-substituted HA (CA) in which 2 to 8 wt% of the material is substituted by carbonate ions, and which also has low concentrations of sodium, magnesium, chlorine and potassium in its crystal structure, resembles the natural mineral phases in bone matrices ^4^. As a result of the similar compositions of the mineral phases in CA and bone, the former is considered to be an ideal clinical biomaterial for bone remodeling.

Osteoconduction at the so-called interface between an implanted biomaterial and bone tissue proceeds via six stages: protein adsorption, osteoblast attachment, osteoblast proliferation, osteoblast differentiation, matrix calcification and bone remodeling ^5^. In the final stage, the osteoclasts play an invaluable reconstructive role by resorbing the osteoid and bioresorbable materials. For this reason, the ability of osteoclasts to resorb HA and CA surfaces and to perform additional remodeling to reconstruct the osteoid into mineralized bone has to be assessed. Even so, although the importance of the interactions between osteoclasts and biomaterials is widely acknowledged, these interactions are not yet fully understood.

There have been many studies evaluating the interfaces between biomaterials and cells, with the aim of controlling cellular behaviours such as adhesion, proliferation and differentiation. Studies of osteoclasts cultured on inorganic biomaterials such as HA, β-TCP and calcium carbonate have been used to evaluate osteoclast activity *in vitro*. Consequently, it has been determined that *in vitro* osteoclast resorption is affected by the type of inorganic materials involved ^6–8^, the incorporation of ions (such as carbonate ^9, 10^ and silicon ^11^), the surface energy ^2^, the physicochemical dissolution process ^12^, the surface roughness of the substrate ^13, 14^ and the surface crystallinity ^15, 16^.

Human osteoclasts adopt a spread cell morphology on A–type CA, in which carbonate ions are substituted at hydroxide sites in the HA crystal structure. This process, which can be demonstrated by actin staining, is caused by the decreased surface energy compared with the values for HA and calcium carbonate ^2^. A previous work reported rabbit osteoclasts resorbed CA but not HA because CA exhibits significant physicochemical dissolution under acidic conditions ^12^. The resorption of CA by human osteoclasts has also been observed in conjunction with AB-type CA crystals based on HA incorporation over 2.4 wt% carbonate ^10^, and mouse osteoclasts have resorbed B–type CA with 7.7 wt% carbonate ^17^. AB–type CA, in which carbonate ions are substituted at hydroxide and phosphate sites in the HA crystal structure, have been found to be resorbed by human osteoclasts along with subsequent acceleration of the proliferation of human osteoblast–like cells and collagen synthesis ^9^. The increased osteoblast proliferation and differentiation associated with osteoclast resorption suggests that CA affects osteoconductivity through the activities of osteoclasts.

Recently, we demonstrated that electrically polarized HA enhances early-stage protein adsorption after *in vivo* implantation ^18^, as well as the initial adhesion and migration of osteoblast-like cells *in vitro* ^19^ and osteoconductivity *in vivo* ^20^, compared with standard HA. Two of the most important factors related to the enhancement of biological reactions through electrical polarization are the attendant increase in the surface free energy and the improved surface wettability of the solid biomaterial ^21, 22^. Other research groups have reported similar effects of electrical polarization on osteoblast-like cells, including greater wettability ^23 24^. Although the promotion of the initial stages of osteoconduction by electrical polarization has been reported, the effects on osteoclast behaviour have not yet been elucidated. We are especially intrigued by the interactions between osteoclasts and polarized biomaterials, which might help to clarify the mechanism by which polarization-induced effects occur in osteoconduction. In the present study, we therefore combined approaches from biology and materials science and used HA and CA with charged surfaces induced by polarization to better understand the interactions between osteoclasts and biomaterials.

## Materials and Methods

### Preparation of biomaterials

HA powder was synthesized from analytical grade calcium hydroxide and phosphoric acid using a wet method ^3^. The resulting HA powder was calcined at 850 °C and then pressed in a mould at 200 MPa, after which the HA compacts were sintered at 1250 °C for 2 h in a saturated water vapor atmosphere. CA was obtained using a modification of the procedure described by Doi et al. ^3, 12^. This CA powder was synthesized from analytical grade calcium nitrate tetrahydrate, disodium hydrogen phosphate and sodium carbonate, with a CO_3_/PO_4_ molar ratio of 5, employing a wet method. CA compacts were produced by pressing this power in a mould at 200 MPa followed by sintering for 2 h in a carbon dioxide atmosphere (to minimize carbonate loss from the CA surface) at 780 °C. Bone slices were also prepared as control samples for osteoclast culture trials, using a procedure previously described in the literature ^3^. In brief, a frozen bovine cortical femur was cut into 130-180 μm thickness slices with a diamond saw (Buehler, Lake Bluff, IL, USA), after which the bone slices were cleaned by ultrasonication in distilled water.

The biomaterial specimens were prepared at thicknesses of 0.8 mm by polishing. After polishing with a 5 μm grain size diamond disk, the HA and CA specimens (ϕ7 mm) were washed in ethanol with ultrasonication. The roughness of each specimen was measured with a colour laser microscope (VK 8500, Keyence, Osaka, Japan) and specimens with an average density of 98% and an Ra value of 0.24 ± 0.05 μm were used for further experimentation.

Some of the HA and CA specimens were electrically polarized using the same process as employed in our previous work ^21^, applying a pair of platinum electrodes at 400 and 350 °C in direct current electric fields of 5 and 2 kV/cm for 1 and 0.5 h, respectively. The electrical polarization provided negatively and positively charged surfaces, and the negatively charged HA and CA surfaces are denoted herein as HA–N and CA–N, respectively, while the positively charged surfaces are referred to as HA–P and CA–P, respectively.

Polarization of the HA and CA specimens was confirmed by thermally stimulated depolarization current (TSDC) measurements. These analyses were carried out in air from room temperature (RT) to 800 °C at a heating rate of 5.0 °C/min according to a method previously described by our group ^22^. The depolarization current was determined with a Hewlett-Packard 4140B pA meter. The stored charge (*Q*)values were calculated from the TSDC spectra according to the equation:

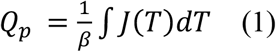

where J(T) is the experimentally determined dissipation current density at temperature T and β is the heating rate. The *Q*, activation energies (*Ea*) and relaxation half-life periods (*τ*) values associated with depolarization were obtained from the TSDC data using the equations:

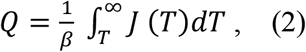

and

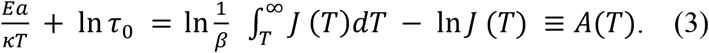

The value of τ, which governs the relaxation process, at 37 °C was calculated using the Arrhenius law:

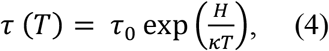

where J(T) is the measured dissipation current density at temperature T, β is the heating rate, τ_0_ is a pre-exponential factor and κ is the Boltzmann constant.

### Surface characterization

X–ray diffraction (XRD) patterns were acquired from HA and CA samples with and without the polarization treatment to assess the presence of various phases, using Cu Kα radiation and operating at 40 kV and 40 mA (Philips PW1700). Attenuated total reflectance FTIR spectroscopy (ATR-FTIR) was also used to examine each polarized HA and CA sample at five different points (Perkin-Elmer spectrum BX spotlight spectrophotometer with a diamond ATR attachment). Prior to acquiring each spectrum, the specimen was held in air at 20 ± 2 °C for 48 h in a desiccator to maintain the same atmosphere as that of the FTIR equipment. Scanning was conducted from 4000 to 400 cm^−1^, with 64 scans averaged for each spectrum.

Surface free energy values for HA and CA were determined based on contact angle measurements using a dual liquid phase method. Specifically, the contact angles of water on the HA and CA surfaces were measured in hydrocarbon oils such as hexane (18.4 mJ⋅m^−2^), heptane (20.1 mJ⋅m^−2^), octane (21.7 mJ⋅m^−2^), decane (23.8 mJ⋅m^−2^) and hexadecane (27.5 mJ⋅m^−2^). The surface free energies were then calculated according to Jouany’s equation:

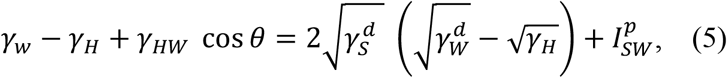

where subscripts W, H and S refer to water, hydrocarbon and solid, respectively, 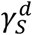 and 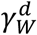 are the dispersion components, and 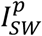 is the nondispersive interaction between the solid and water as expressed by:

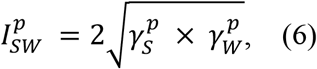

where 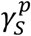 and 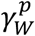 are the polar (nondispersive) components.

According to Fowkes, the work of adhesion (W) between a solid and water can be divided into two interaction components. These represent dispersive and nondispersive interactions ^25^ and can be written as:

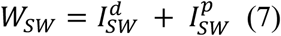

and

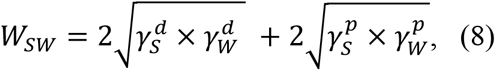

where the dispersive component for water 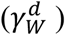 has a value of 21.8 and the polar component for water 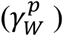 has a value of 51.0 ^25^. The geometric mean expression used for 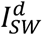 above is based on the work of Fowkes, while that for 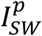 is an extended one.

The surface wettability of each material was assessed by performing contact angle measurements in air (Kyowa Interface Science, DropMaster DM-500) with distilled, deionized water (Merck Millipore, Direct-QUV). These measurements were performed using as-polarized samples and samples one month after the polarization treatment. The contact angles were calculated using Young’s equation:

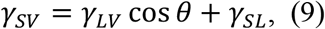

where the subscripts S, L and V refer to the solid, liquid and vapor phases, respectively.

### Human osteoclast cultures

Peripheral mononuclear blood cells (PBMCs) were obtained from blood donated by healthy human males, as previously described ^3, 26^. The donation protocol was approved by the Human Subjects Committee of the University of Turku and Tokyo Medical and Dental University. Briefly, anticoagulated human blood with heparin was diluted 1:1 (v/v) with phosphate buffered saline (PBS) and buffy coats layered over Ficoll-Paque Plus solution (Amersham Pharmacia Biotech, Uppsala, Sweden) were diluted 1:1 (v/v) with PBS and centrifuged at 1500 rpm for 15 min. The cell layer above the Ficoll-Paque Plus was collected and washed twice with PBS and resuspended in α-MEM including 10% FBS, 100 IU/mL penicillin and 100 μg/mL streptomycin. The enriched PBMCs were seeded onto the specimens at a density of 1 × 10^6^ cells/cm^2^ and cultured in a humidified atmosphere containing 5% CO_2_ at 37 °C for 60 min. The specimens with the attached PBMCs were subsequently placed into 24 well plates and then differentiated by the addition of 20 ng/mL RANKL (Peprotech 310-01) and 10 ng/mL M-CSF (R&D, 216-MC) in α-MEM including 10% FBS, 100 IU/mL penicillin and 100 μg/mL streptomycin. Half of the media in each sample were changed every three to four days. Cells were cultured on the specimens for 14 days, washed twice with PBS, and then fixed with 4% paraformaldehyde for 20 min.

### Staining

To confirm the differentiation of the PBMCs into osteoclasts, the cells adhering to the specimens were stained for TRAP (Sigma, No. 387) and the number of TRAP positive multinucleated cells per specimen was counted on each surface. A total of at least 30 fields on each specimen was measured to obtain an average. Cell morphologies were visualized by blocking the cells with 5% goat serum in a solution of 0.1% Triton X-100 in PBS. A mouse monoclonal anti-vinculin antibody in the blocking solution was added for 1 h at RT. Following extensive washing with PBS, the specimens were incubated in Alexa-conjugated goat anti-mouse immunoglobulin in a blocking solution containing rhodamine phalloidin for 1 h at RT. After Hoechst staining, fluorescent emissions were observed using a fluorescence microscope (Olympus IX71). The diameters and thicknesses of the actin rings were measured at random to quantify the adhesion of the cells, using <u>NIH image®</u>. These measurements were performed for a minimum of 50 cells on each specimen to obtain an average, and the quantity of nuclei per cell was calculated to quantify the fusion of the cells. At least 50 cells were assessed on each specimen to obtain an average.

### Analysis of resorption pits

After each specimen was examined using a microscope, the osteoclasts were removed from the specimens by scrubbing with a brush, following which each sample was washed with distilled water and dried. The resorption activities of the osteoclasts were assessed by examining the morphological parameters of the resorption pits. The dried specimens were examined with a colour laser microscope (VK8500, Keyence, Osaka, Japan) equipped with an image analysis system, and the depth of each pit was measured to quantify the amount of osteoclast resorption. The measurements were performed for a minimum of 30 resorption pits on each specimen to obtain an average.

### Statistical analysis

Accurate quantifications of the different samples were achieved by performing more than three independent experiments. Statistical analysis between groups was performed by an analysis of variance (ANOVA) with the Tukey formula for *post hoc* multiple comparisons, using the SPSS software package (version 22, Chicago, IL). A statistical significance level of *p* < 0.05 was used for all tests. All data are expressed herein as mean ± standard deviation (SD).

## Results

Typical TSDC patterns for the polarized HA and CA are shown in Figs. 1a and 1b, respectively. The TSDC curves for the HA and CA respectively increased at approximately 200 and 400 °C, reached maxima at approximately 510 and 580 °C, and then gradually decreased. The maximum current density was approximately 10 nA/cm^2^ for HA and 10,000 nA/cm^2^ for CA. The *Q* calculated from the TSDC data was 26 μC/cm^2^ for HA and 13 mC/cm^2^ for CA. The *Ea* obtained from the curves were 0.9 and 1.2 eV for HA and CA, respectively, while the *τ* at 37 °C were calculated to be 1 × 10^3^ and 2 × 10^15^ years. These differences in the stored charges, activation energies and half periods are attributed to the different charged carrier particles participating in the polarization process. These comprised protons for HA and oxygen ions for CA ^27^.

**Fig. 1.**
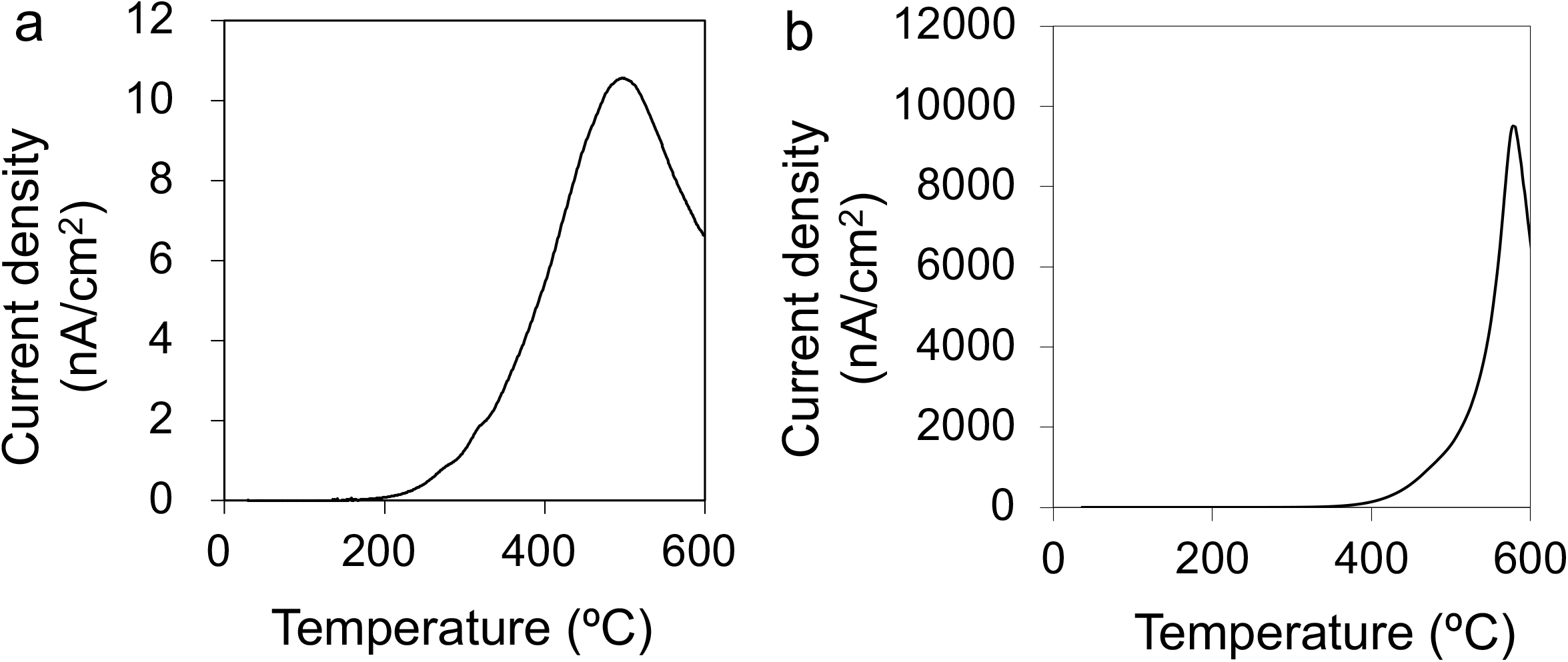
Typical TSDC curves obtained from (a) HA and (b) CA samples after polarization treatments.

The XRD patterns obtained from the HA and CA samples both with and without the polarization treatment were equivalent to the HA standard pattern (ICDD No.9–432), meaning that each surface consisted of a single hexagonal HA phase (Fig. 2). However, the peak related to the (002) diffraction for CA appeared at a lower angle than that for HA, suggesting that carbonate ions had been substituted at some of the phosphate ion sites ^12^. The ATR-FTIR spectra of all specimens contained peaks assigned to phosphate ions at 1045, 1089, 601, 575 and 567 cm^−1^ (Fig. 3), and peaks attributed to stretching and bending vibrations of hydroxide ions were clearly observed at 3570 and 630 cm^−1^ in the HA spectra. The spectra obtained from CA also contained carbonate peaks at 1415, 1450 and 871 cm^−1^. These data confirm that the sintered CA specimens used in this study comprised B-type CA containing approximately 8 wt% carbonate ions substituted at phosphate sites in the HA lattice ^17^.

**Fig. 2.**
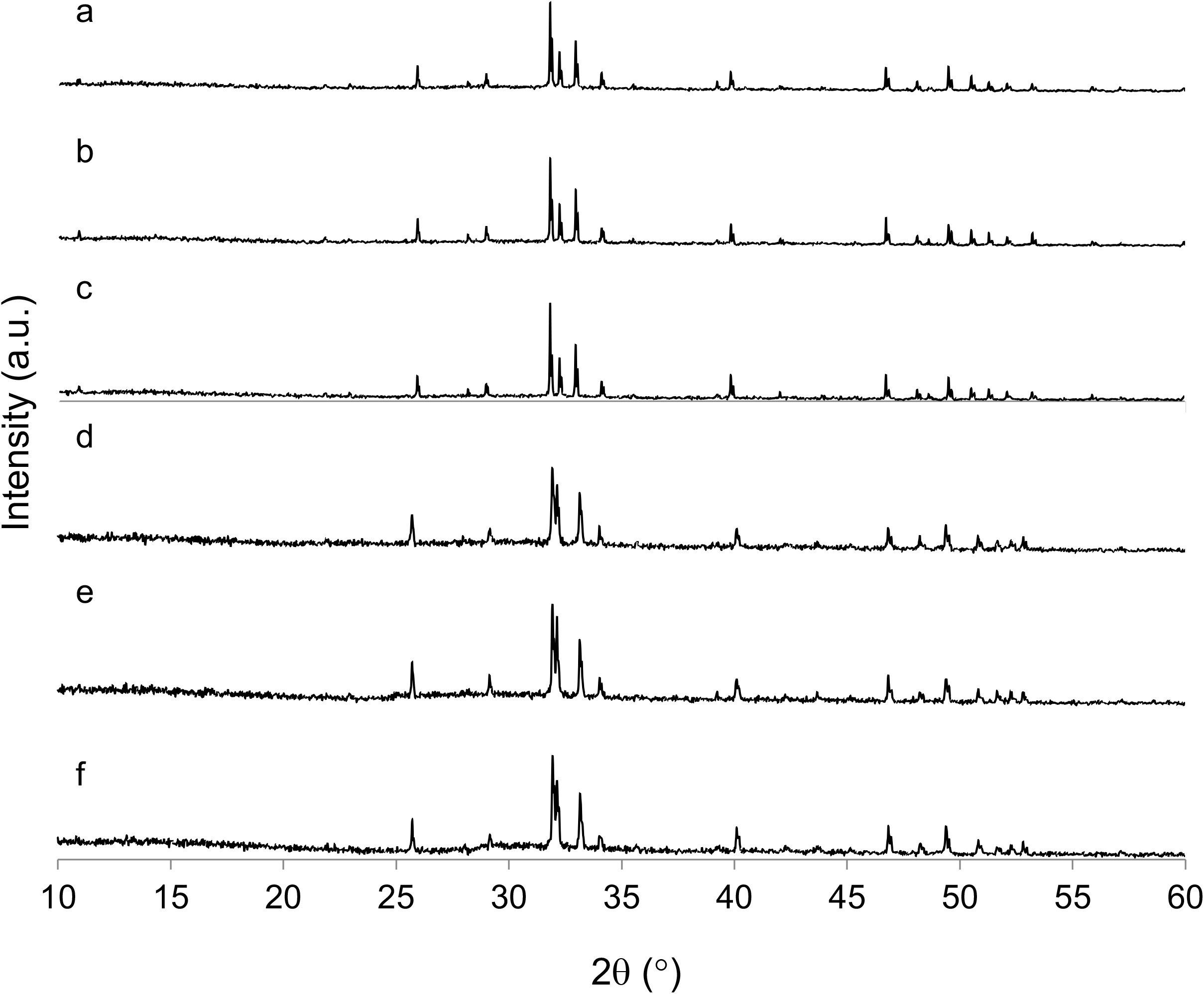
XRD patterns obtained from (a) standard HA, (b) HA–N, (c) HA–P, (d) standard CA, (e) CA– N and (f) CA–P. Note that each is extremely similar to that of single phase hexagonal HA.

**Fig. 3.**
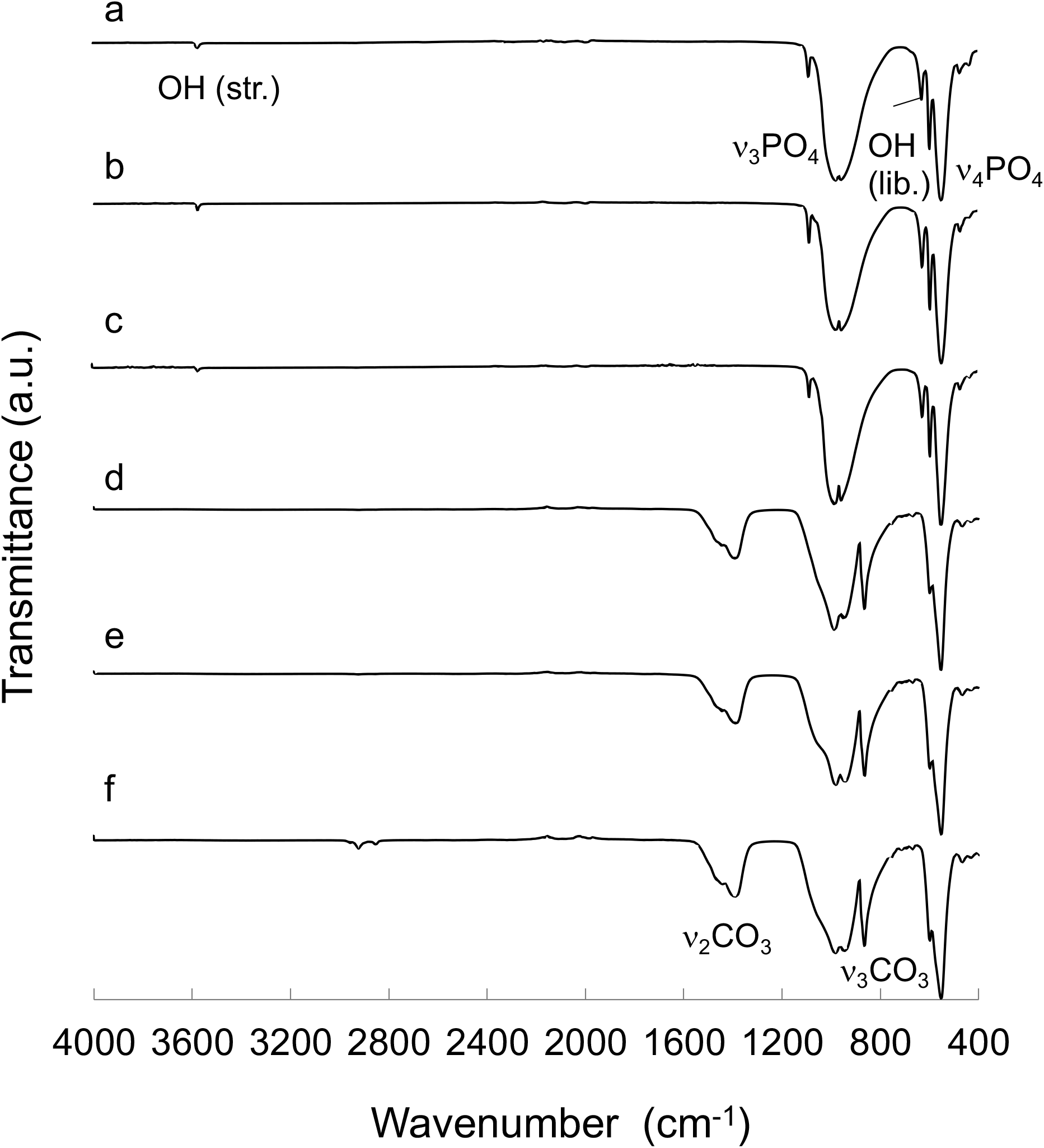
ATR-FTIR spectra obtained from (a) standard HA, (b) HA–N, (c) HA–P, (d) standard CA, (e) CA–N and (f) CA–P.

The contact angles in hydrocarbon oils determined for HA and CA specimens with and without polarization using the dual liquid method are presented in Fig. 4a. The dispersive and polar components of the surface free energy values were calculated from the slopes and y-intercepts of these plots, and the surface free energies were obtained by summing the dispersion and polar components (Table 1). The resulting surface energies were 38.6 mJ/m^2^ for standard HA and 51.1 mJ/m^2^ for standard CA. The surface energy values for initial polarized HA and CA were found to be increased by factors of approximately 1.7 and 1.5 times, respectively, relative to the unpolarized samples. After one month, these standard values for HA and CA were found to increase to 40.3 and 55.0 mJ/m^2^, respectively. In addition, the surface energy values for the polarized HA and CA after one month were found to be increased by factors of approximately 1.7 and 1.5 times, respectively, relative to the unpolarized of one-month aged samples.

**Fig. 4.**
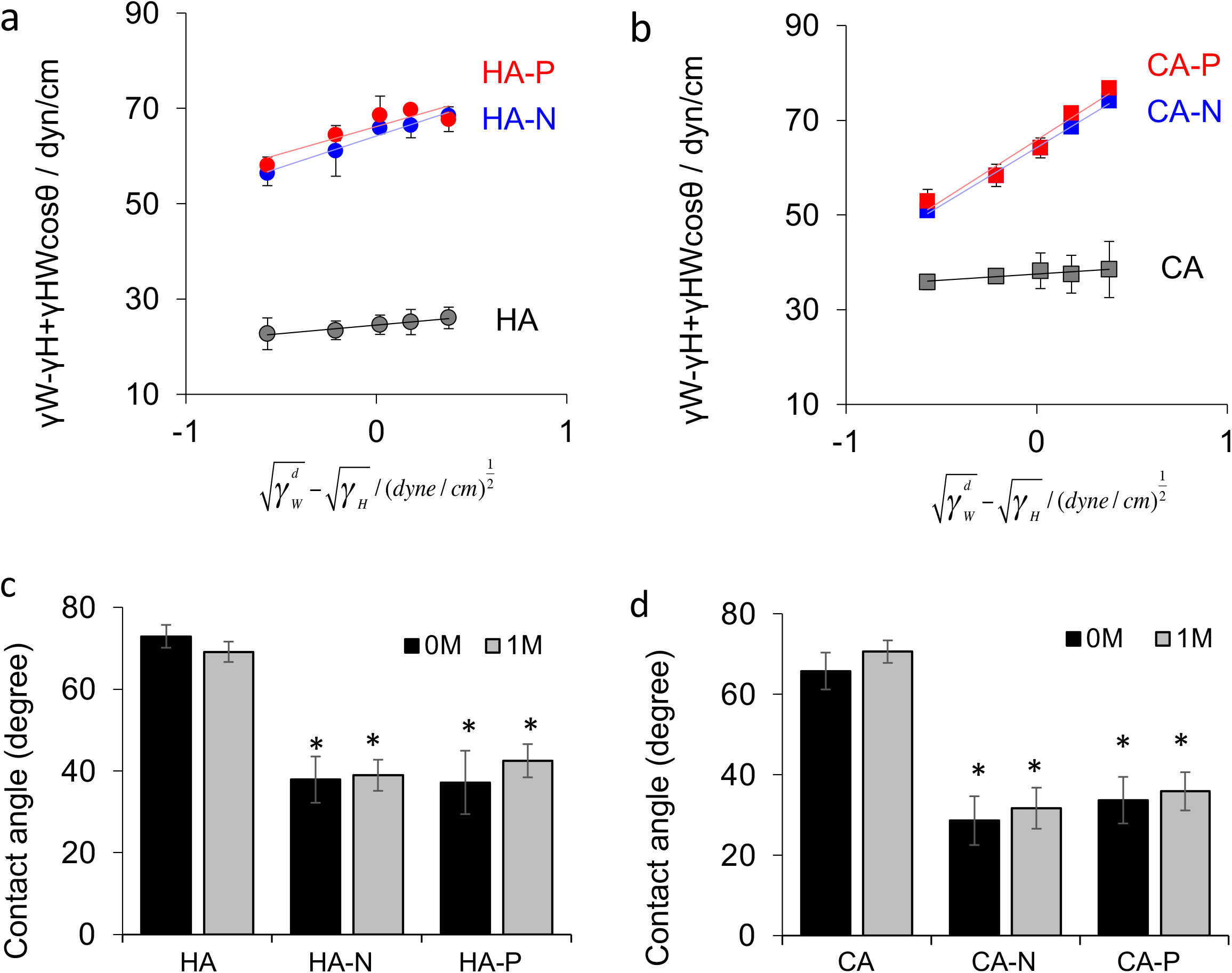
Contact angles determined using the two phase liquid method in hexane, heptane, octane, decane and hexadecane with distilled, deionized water for (a) HA and (b) CA with and without polarization treatments. Black, blue and red symbols indicate standard, negatively charged and positively charged surfaces, respectively. Contact angle data for (c) HA and (d) CA with and without polarization treatments determined using distilled, deionized water in the as-polarized state and one month after the polarization treatments. The HA and CA surfaces presented higher angles, indicating that these materials were more hydrophobic.

**Table 1.** Surface free energies of HA and CA samples with and without polarization treatments as calculated using Jouany’s equation. The dispersion and polar components of the surface free energies were calculated based on the slopes and y-intercepts of plots of contact angles determined in hydrocarbon oils (Figs. 4a and 4b). The surface free energy values were calculated by summing the dispersion and polar components.

|  | As-polarized |  |  | After 1 month |  |  |
| --- | --- | --- | --- | --- | --- | --- |
| | Dispersive components<br>: $\gamma_s^d$<br>(mJ/m<sup>2</sup>) | Polar components<br>: $\gamma_s^p$<br>(mJ/m<sup>2</sup>) | Surface free energy<br>(mJ/m<sup>2</sup>) | Dispersive components<br>: $\gamma_s^d$<br>(mJ/m<sup>2</sup>) | Polar components<br>: $\gamma_s^p$<br>(mJ/m<sup>2</sup>) | Surface free energy<br>(mJ/m<sup>2</sup>) |
| HA | 2.23 | 36.4 | 38.6 | 2.2 | 38.1 | 40.3 |
| HA-N | 11.8 | 55.4 | 67.2 | 17.0 | 51.6 | 68.6 |
| HA-P | 13.3 | 54.8 | 68.1 | 11.0 | 53.6 | 64.6 |
| CA | 6.10 | 45.0 | 51.1 | 9.84 | 45.2 | 55.0 |
| CA-N | 17.7 | 62.0 | 79.7 | 17.6 | 55.3 | 72.9 |
| CA-P | 13.6 | 61.2 | 74.8 | 14.8 | 57.3 | 72.1 |

The contact angle values determined in air using distilled, deionized water for HA and CA were significantly decreased after the electrical polarization treatment (Figs. 4c and 4d). The angles were 73 ± 2.8° for the standard HA and 66 ± 5.1° for the standard CA prior to polarization while the values were 37 ± 4.9°, 33 ± 5.6°, 34 ± 5.8° and 29 ± 6.1° for the HA–N, HA–P, CA–N and CA–P, respectively, after polarization. The surfaces of the polarized HA and CA therefore exhibited lower contact angles, meaning that the wettability of each negative and positive surface was improved. The contact angle values were remeasured one month after the polarization treatment and compared with those of the as-polarized surfaces and no significant differences were found. Thus, the increased surface free energy and improved surface wettability were maintained for at least this length of time.

Osteoclasts derived from human PBMCs were positively stained for TRAP on the surfaces of all specimens after culturing with RANKL and M-CSF for 14 days (Fig. 5a). The TRAP staining showed that the PBMCs adhering to the specimens had differentiated into osteoclast precursors or osteoclasts. Some TRAP-positive cells adhering to the specimens were small and mononuclear, suggesting that they had not completely differentiated into osteoclasts. The quantities of multinuclear TRAP-positive cells (which indicate osteoclast differentiation) were significantly larger on the CA samples compared with the numbers on the HA samples (Fig. 5b), while the bone slices exhibited the highest concentration of multinuclear TRAP-positive cells.

**Fig. 5.**
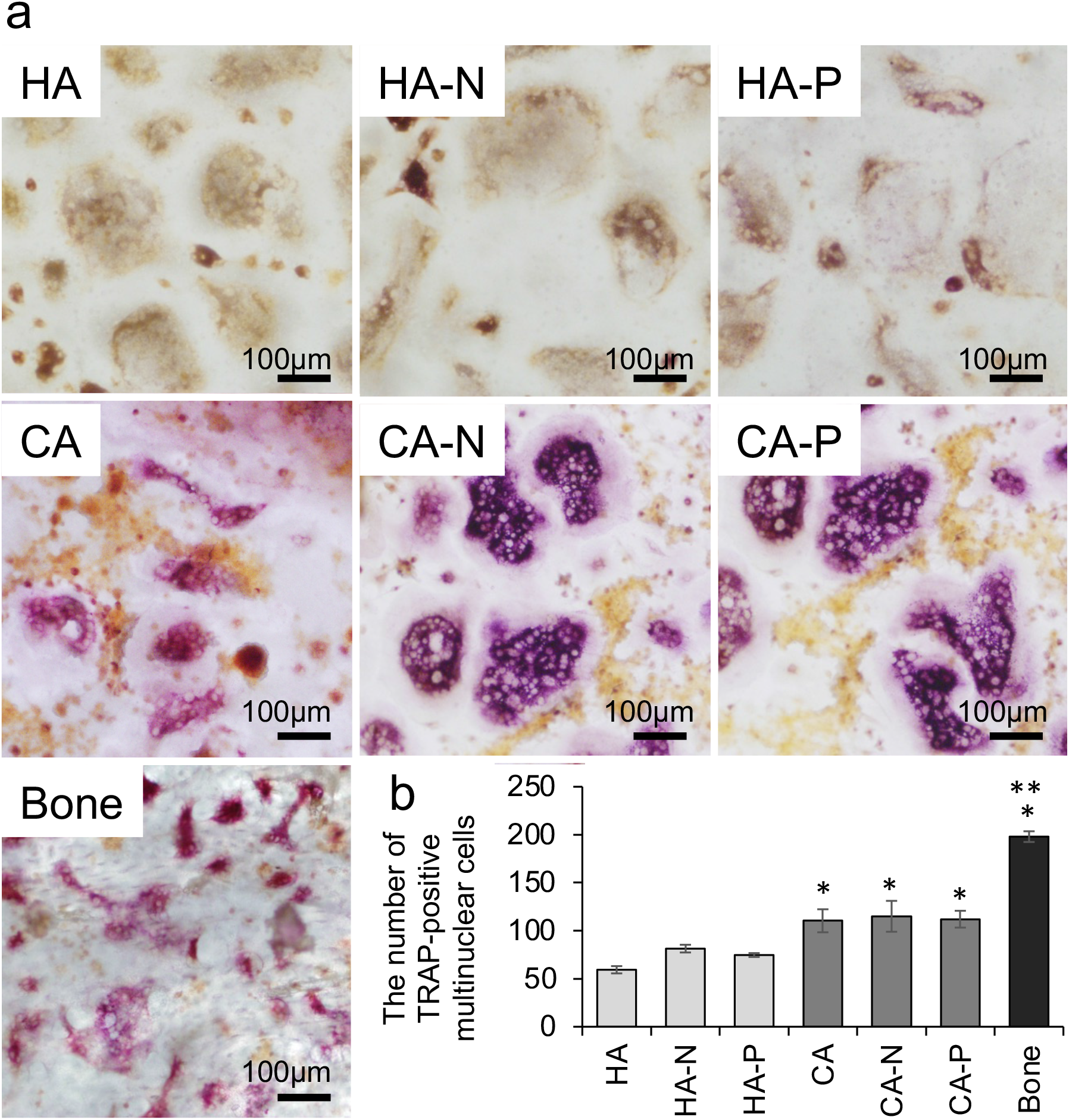
(a) TRAP staining of the cells cultured on bone slices and on HA and CA samples with and without polarization treatments. The bars indicate 100 μm. (b) Quantities of multinuclear TRAP-positive cells used to quantify the differentiation of PMBCs into osteoclast precursors or osteoclasts on seven specimens. The number of multinuclear TRAP-positive cells was significantly increased on the CA samples and bone slices compared to the HA specimens. *p<0.002, **p<0.001 compared with the others. Error bars indicate ± one standard deviation.

Figure 6 shows the results of vinculin (green), actin (red) and nuclei (blue) labelling of the adhered cells derived from human PBMCs (Fig. 6a). On each surface, osteoclast precursor cells adhered and differentiated into giant multinucleated cells in the induction medium. Actin rings, which are specific markers for activated osteoclasts, were also formed in the osteoclasts on the HA, CA and bone slices. The osteoclast morphologies were quantified by assessing the number of nuclei in each cell, the thicknesses of the actin rings and the diameters of the actin rings. The number of nuclei in each cell was essentially equivalent for all specimens, showing that the substrate materials did not affect cell fusion during the osteoclast differentiation process (Fig. 6b).

**Fig. 6.**
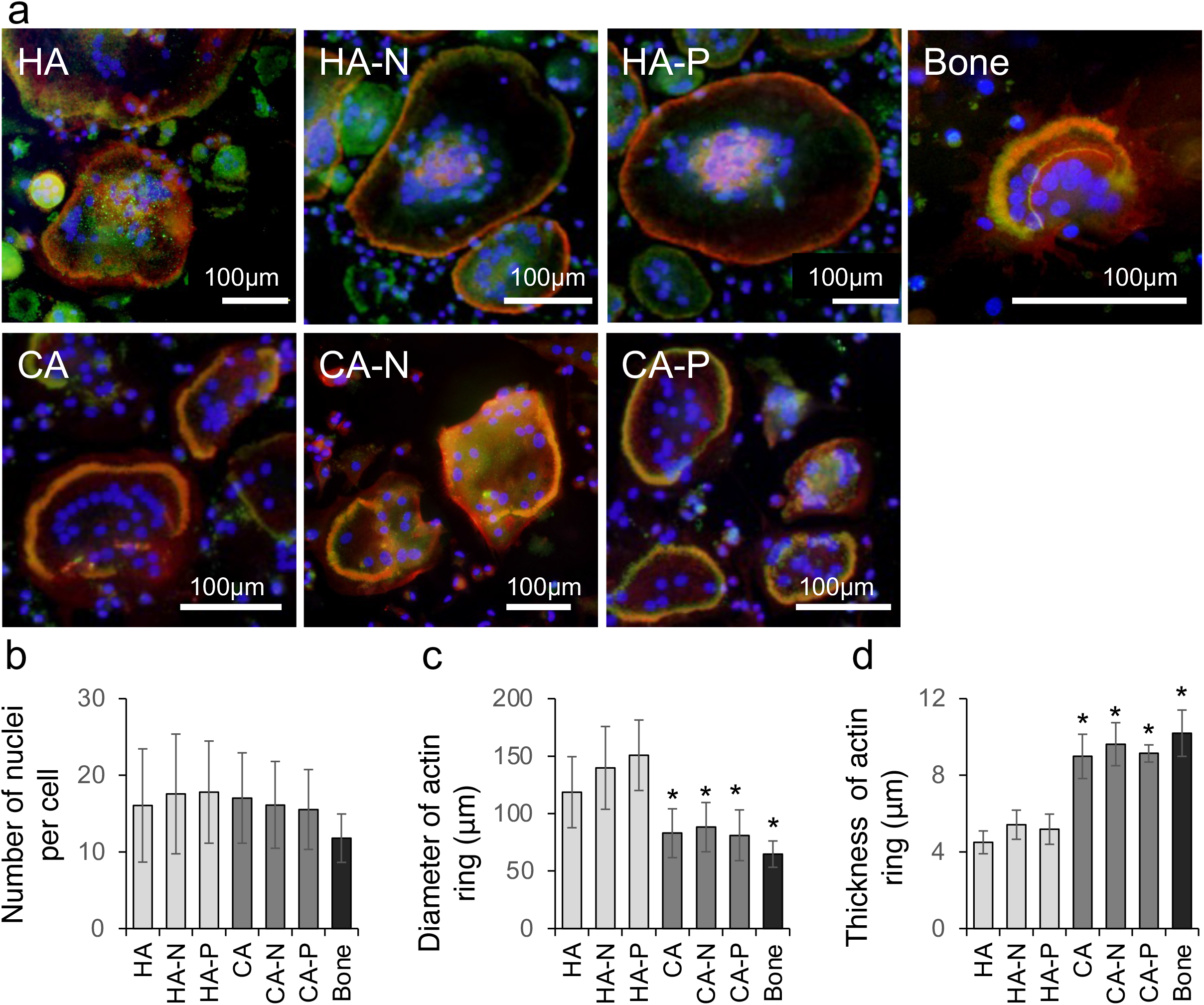
(a) Morphologies of the osteoclasts adhering to bone slices and to HA and CA samples with and without polarization treatments, as visualized by the fluorescent staining of actin filaments (red), vinculin (green) and nuclei (blue). Actin ring structures were formed in the osteoclasts on each surface and the vinculin was co-localized with the actin rings in the cells cultured on the bone slices and CA specimens, while the vinculin was scattered throughout the cytoplasm of the cells on the HA samples. The bars indicate 100 μm. (b) Quantities of nuclei in each osteoclast as a means of quantifying differences in osteoclast morphology. The number of nuclei in each cell was almost the same for all specimens. (c) The diameters of actin rings in osteoclasts as a means of quantifying the differences in the actin ring morphologies. The actin rings in the osteoclasts cultured on the CA samples and bone slices were significantly smaller than those on the HA specimens. (d) The thicknesses of actin rings in osteoclasts as a means of quantifying the differences in actin ring morphologies. The actin rings in osteoclasts cultured on the CA samples were thicker and similar to those on the bone slices, whereas the actins rings of osteoclasts cultured on the HA samples were thin. *p<0.001 compared with the others. Error bars indicate ± one standard deviation.

Three differences were identified in the fluorescence images obtained from the HA and CA specimens, one of which was the vinculin-immunoreactive distribution (Fig. 6a). Specifically, vinculin was co-localized with the actin rings of the osteoclasts cultured on the bone slices and CA but scattered throughout the central regions of the osteoclasts cultured on the HA. The second difference was in the actin ring sizes of the osteoclasts (Fig. 6c), such that the ring sizes of osteoclasts cultured on the CA and bone slices were significantly decreased compared to those on the HA. These larger actin rings indicated that the spread of osteoclasts was enhanced on the latter material. The third difference was the thickness of the osteoclast actin rings (Fig. 6d). The rings of osteoclasts cultured on the CA samples were thicker and similar to those seen on the bone slices, whereas those of the osteoclasts cultured on the HA specimens were thin.

Resorption pits were observed on the bone slices and on the HA and CA surfaces using three– dimensional laser microscopy (Fig. 7a). However, distinct pits were formed on the bone slices and CA samples, whereas ambiguous, shallow pits appeared on the HA sample. The resorption pits formed by osteoclasts on the CA samples were also approximately 11 times deeper than those on the HA sample (Fig. 7b). The pit depths on the CA–N and CA–P samples were approximately 18 and 21 times greater than those on the standard HA sample and 1.6 and 1.9 times greater than those on the standard CA sample, respectively.

**Fig. 7.**
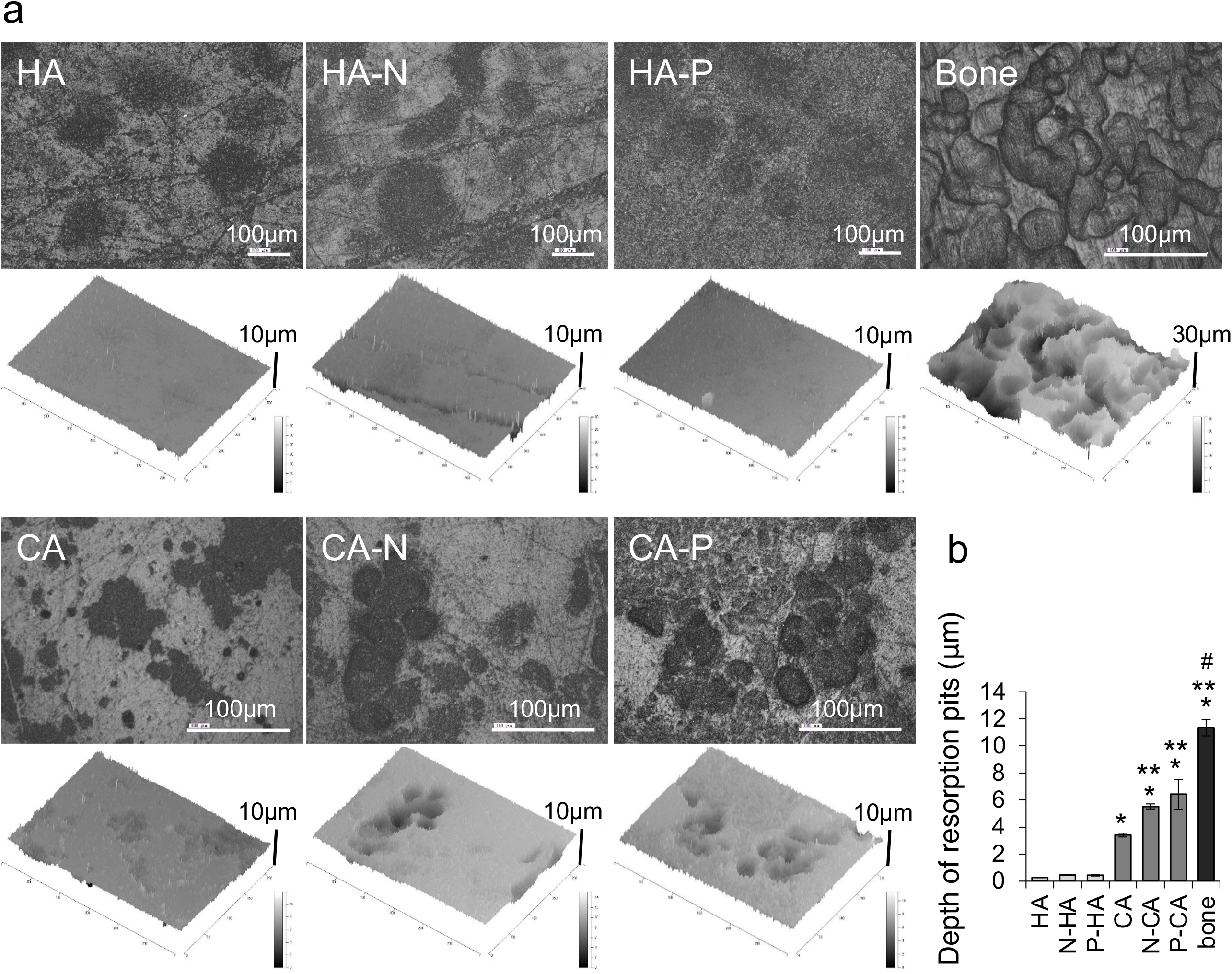
(a) Images of the resorption pits formed on bone slices and on HA and CA samples with and without polarization treatments. The bars indicate 100 μm. (b) The depths of the surfaces after osteoclast culturing and cell scrubbing as a means of quantifying the differences in resorption pits. The pits formed by the osteoclasts on bone slices were deeper than those on the other specimens while the pits on the CA samples were larger than those on the HA samples. The pits on the polarized CA were also deeper than those on the standard CA surface. *p<0.001, **p<0.002, #p< 0.001 compared with the others. Error bars indicate ± one standard deviation.

## Discussion

The results of this work show that the concentration of carbonate ions in an apatite crystal structure affects the induction of osteoclastogenesis, and that polarization modifies the resorption of human osteoclasts. Here, we discuss the effects of the carbonate ions and surface free energy in relation to the conventional understanding of surface characteristics and cell activities.

The trials involving osteoclast cultures showed that the human PBMCs differentiated into mature osteoclasts and were activated to resorb the substrate when using the CA samples. These results are in agreement with our previous study ^3, 17^, in which standard CA was found to enhance osteoclast differentiation, as indicated by increases in the number of TRAP-positive multinuclear cells. This prior work also indicated the formation of thick, compact actin rings that create tight seals between the osteoclasts and the CA, as well as the formation of deeper resorption pits compared with those formed on standard HA. The characterization of CA specimens using XRD and ATR-FTIR revealed that the CA surfaces consisted of a single, highly crystalline HA phase and that phosphate ions were partially substituted for carbonate ions, meaning that the material could be classified as B– type CA ^17^ and had properties similar to bone. It should be noted that the mineral component of a bone matrix is nonstoichiometric HA with partial substitution of carbonate, sodium, magnesium, chlorine and potassium ions within its crystal structure ^4^. Therefore, one possible explanation for the variations in osteoclast differentiation and resorption is that changes in the material properties with the substitution of carbonate ions into the HA crystal structure increased the similarity to bone tissue.

Osteoclast activity is an essential aspect of controlling bone remodeling and is related to the organization of the actin cytoskeleton to form a sealing zone that anchors the osteoclasts to the bone matrix. The morphology of the actin rings is affected by surface roughness ^13, 14^, crystallinity of the solid surface ^15^, solubility ^28^ and surface free energy ^29^. In the present study, the synthesized substrates had approximately equivalent surface roughness values as a result of polishing. In addition, the XRD data showed that the material surfaces consisted of a single highly crystalline HA phase. Therefore, the solubilities and surface energies of the synthesized specimens are believed to have had the greatest effects on osteoclast resorption.

The higher solubility of CA would be expected to promote osteoclast resorption by generating a sealing zone between the osteoclasts and the bone matrix. Calcium and phosphate ions are released from the bone matrix during osteoclast resorption, and this release stimulates the resorption of surrounding osteoclasts. The sensitivity of osteoclasts to extracellular calcium concentrations is reported to vary throughout the osteoclast activation process, which comprises resting, migrating and resorbing ^30^. Resorbing osteoclasts are not sensitive to increased extracellular calcium concentrations, while resting osteoclasts are more sensitive. Specifically, resting osteoclasts are stimulated by calcium ions released from the bone matrix and subsequently differentiate to resorb the intact areas of the bone matrix. Activated osteoclasts on CA will resorb the CA surface to release calcium ions ^31^ and phosphate ions ^32^ into the cell culture medium. These ions then stimulate and differentiate the surrounding resting osteoclasts such that the intact CA surface is resorbed.

Another possible explanation for the observed osteoclast formation on the CA is acidification resulting from the release of carbon dioxide or bicarbonate from the CA surface. These species could possibly be released along with calcium and phosphate ions under acidic conditions, such as in the case of surgical wounds or injuries. The acidic conditions at these sites are known to induce osteoclast resorption ^33^. In the present study, the osteoclasts displayed a more spread cell morphology on the HA samples compared to that on the CA specimens (Figs. 6a and 6c) and formed deeper resorption pits on the surfaces of the CA materials (Fig. 7). These results can possibly be attributed to the higher solubility of CA because the standard CA exhibited increased solubility compared to the standard HA^17^.

The surface energy value is known to affect the initial adherence, spread and formation of the collagen fibrils of human osteoblast-like cells ^29^ as well as the cellular morphology of human osteoclasts ^2^. Human osteoclasts cultured on A–type CA (in which carbonate ions are substituted for hydroxyl sites in the HA lattice) displayed a more spread cell morphology, as shown by actin staining ^2^. Redey et al. have suggested that the different cell morphology on A–type CA results from a decrease in the polar component of the surface free energy compared with HA. Such reports suggest that the surface free energy affects cell activities on CA. Although the ion substitution sites in the crystal structures were different between A–type CA and B–type CA, the B–type CA used in the present study showed an increase in surface energy by a factor of approximately 1.5 following polarization (Fig. 4). In addition, the polarization treatment accelerated osteoclast resorption on the CA (Fig. 7). Overall, these results indicate that increases in the surface free energy of the polarized CA promoted osteoclast resorption.

Another possible explanation for the accelerated osteoclast resorption on the polarized CA is based on changes in protein conformation. Many different proteins that are involved in cell adhesion were included in the cell culture medium, and increasing the surface free energy could have changed the conformation of the proteins related to osteoclast adhesion. Each CA specimen was immersed in the cell culture medium prior to osteoclast culturing, and so was presumably covered with a similar amount of adsorbed proteins. Taking into account both the evident changes in surface free energy caused by ion substitution in the apatite crystal structure ^29^ and variations in the conformations of adhesion proteins as a consequence of surface chemistry ^34, 35^, it is possible that the conformation of proteins adsorbed on the substrates varied as a consequence of differences in the surface free energy values of the samples. Variations in the conformations of the adhesion proteins could possibly lead to differences in the exposed domains of the proteins adsorbed on the specimens.

During bone remodeling, the osteoclast resorption period is known to be 30 to 40 days and is followed by bone formation over a period of 150 days ^36^. In the work reported herein, the polarization effects were evidently maintained over at least one month because the increased surface free energy and improved surface wettability were retained over this time span (Table 1 and Figs. 4c and 4d), and because the surface charges on the HA induced by polarization were maintained during sterilization and cell culture ^37^. Considering that the τ values at 37 °C obtained from the TSDC curves were 1 × 10^3^ years for HA and 2 × 10^15^ years for CA (Fig. 1), the surface charges would be expected to remain throughout the bone remodeling period. This stability of the surface charges indicates that electrical stimulation and the concurrent polarization could be beneficial as a clinical treatment to accelerate bone remodeling. The results of this work should contribute to the future design of cell mediated bioresorbable biomaterials capable of resorption by osteoclasts and of serving as scaffolds for bone regeneration.

## Conclusions

The results of the present study provide new and important information regarding the surface characteristics of polarized ceramic biomaterials and the behaviour of osteoclasts on these materials. Polarization was found to improve the surface wettability of HA and CA as a result of increases in surface free energy, and this effect was still present after one month. In addition, trials in which osteoclasts were cultured on various substrates showed that polarized CA accelerated osteoclast resorption but did not affect the TRAP staining and morphology of osteoclasts.

## Acknowledgements

We thank Ms. Naoko Hori and Ms. Takako Takuma for their technical assistance. This study was financially supported by the Turku Collegium for Science and Medicine, a Grant–in–Aid for the Promotion of Joint International Research (Fostering Joint International Research) (no. 15KK0299), Grants–in–Aid for Scientific Research (C) (nos. 17K10957 and 20K09454) and the Murata Science Foundation.

## Notes

### Competing Interest Statement

The authors have declared no competing interest.

